# The intestinal microbiota programs diurnal rhythms in host metabolism through histone deacetylase 3

**DOI:** 10.1101/580613

**Authors:** Zheng Kuang, Yuhao Wang, Yun Li, Cunqi Ye, Kelly A. Ruhn, Cassie L. Behrendt, Eric N. Olson, Lora V. Hooper

## Abstract

Circadian rhythmicity is a defining feature of mammalian metabolism that synchronizes metabolic processes to day-night light cycles. Here, we show that the intestinal microbiota programs diurnal metabolic rhythms in the mouse small intestine through histone deacetylase 3 (HDAC3). The microbiota induced expression of intestinal epithelial HDAC3, which was recruited rhythmically to chromatin and produced synchronized diurnal oscillations in histone acetylation, metabolic gene expression, and nutrient uptake. HDAC3 also functioned non-canonically to coactivate estrogen related receptor α (ERRα), inducing microbiota-dependent rhythmic transcription of the lipid transporter gene *Cd36* and promoting lipid absorption and diet-and jet lag-induced obesity. Our findings reveal that HDAC3 integrates microbial and circadian cues to regulate diurnal metabolic rhythms, and pinpoint a key mechanism by which the microbiota controls host metabolism.

**One sentence summary:** The intestinal microbiota induces daily metabolic rhythms and controls lipid uptake through the enzyme histone deacetylase 3.

---

Mammalian metabolism is acutely sensitive to environmental cues. These include circadian day-night light cycles, which govern when food is consumed, and the microbiome, which impacts how food is digested. The intestinal microbiota shapes the expression of host metabolic pathways and influences the development of obesity (*1-4*). Environmental light signals entrain rhythms in gene expression that are synchronized with the day-night light cycle through the circadian clock (*5-7*). This circadian synchronization is fundamental to metabolic processes that must be coupled to diurnal sleep-wake and feeding-fasting cycles. It is becoming clear that there is cross-talk between the microbiota and circadian pathways that impacts host metabolism (*5-9*). However, the molecular mechanisms by which diurnal light signals and the microbiota converge to regulate metabolism are largely unknown.

A key mechanism by which the circadian clock generates rhythms in metabolic gene expression is by regulating the recruitment of histone modifiers to chromatin (*10-13*). We therefore investigated whether the microbiota might regulate host circadian gene expression by altering histone modifications. We collected small intestinal epithelial cells (IECs) from conventional (CV) and germ-free (GF) mice across a 24-hour cycle (fig. S1A). We then performed RNA sequencing (RNA-seq) and chromatin immunoprecipitation followed by sequencing (ChIP-seq) of H3K9ac and H3K27ac, two histone acetylation marks that indicate active promoters and enhancers. As expected, both acetylation marks localized to the regulatory regions of actively transcribed genes (fig. S1B,C). Both H3K9ac and H3K27ac showed synchronized diurnal oscillations in CV IECs with peaks between Zeitgeber time (ZT) 8 and 16 (where ZT0 is light on and ZT12 is light off), and troughs between ZT20 and ZT4 (Fig. 1A,B).

**Figure 1:**
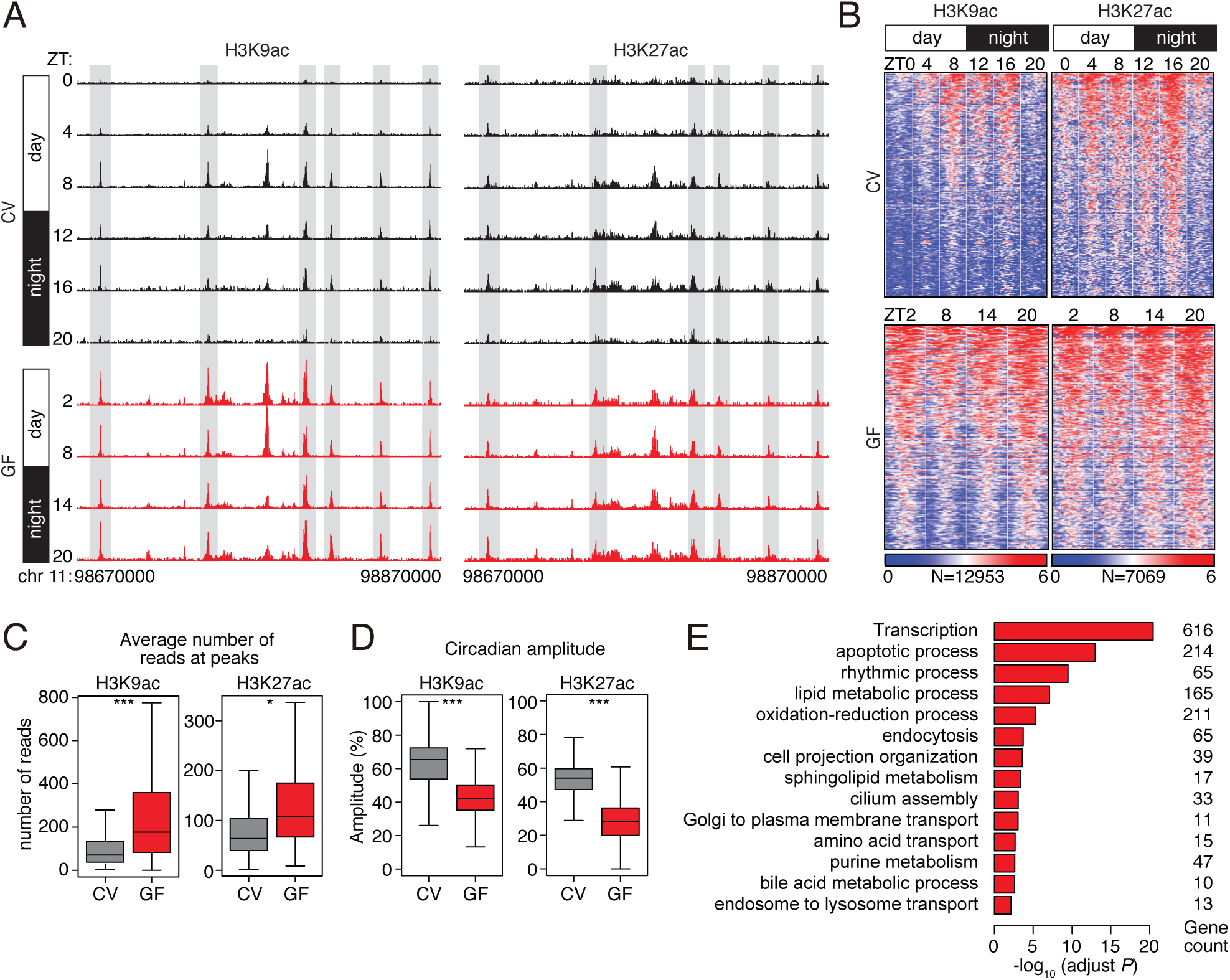
Histone acetylation in small intestinal epithelial cells exhibits synchronized diurnal rhythmicity that depends on the microbiota. **(A)** Genome browser view of the 200 kb region surrounding the *Nr1d1* locus, showing ChIP-seq analysis of H3K9ac and H3K27ac marks in small intestinal epithelial cells. The analysis was done across a circadian cycle in CV and GF mice. Each track represents the normalized ChIP-seq read coverage at a single time point. Examples of peaks showing microbiota-dependent diurnal rhythmicity are highlighted in gray. N=3 pooled biological replicates per library. **(B)** Heat maps of H3K9ac and H3K27ac signals (log (reads at 50 bp windows)) from −1 to +1 kb surrounding the centers of all cycling H3K9ac and H3K27ac peaks (adjust *P*<0.01 by JTK). Each peak in the genome is represented as a horizontal line, ordered vertically by signal strength, and the analysis was done across a circadian cycle in CV and GF mice. The number of peaks in the genome is indicated at the bottom. The blue-red gradient indicates the coverage or signal strength (normalized uniquely mapped reads per 20 million reads). Intensity **(C)** and amplitude **(D)** of H3K9ac and H3K27ac peaks in CV and GF mice. \**P*<0.05, \*\*\**P*<0.001 by one-tailed paired *t*-test. Means±SEM (error bars) are plotted. **(E)** Enriched GO categories in genes having H3K9ac-and H3K27ac-targeted genes as determined by the DAVID GO analysis tool. ZT, Zeitgeber; CV, conventional; GF, germ-free.

Although the majority of the H3K9ac and H3K27ac marks were present in both CV and GF mice (fig. S1D), most of the peaks showed markedly dampened oscillations in GF mice (Fig. 1A-D). Average signal intensities of both marks were increased and circadian amplitudes were decreased in GF mice (Fig. 1C,D). Genes involved in metabolic processes including nutrient transport and lipid metabolism were enriched near cycling histone acetylation peaks (Fig. 1E), and coincided with dampened oscillations in the abundances of transcripts encoding proteins involved in nutrient transport and lipid metabolism (fig. S2A-C). Thus, the microbiota drives synchronized diurnal oscillations in histone acetylation in the small intestine, and many of the targeted genes regulate nutrient uptake and metabolism. This behavior is distinct from the effect of the microbiota on histone acetylation in the colon, where the overall genome-wide rhythmicity of acetylation is not lost after antibiotic depletion of the microbiota (*5*).

H3K9ac and H3K27ac marks cycled diurnally in CV mice but remained constitutively high in GF mice, suggesting a possible mechanism involving a histone deacetylase (HDAC) that is expressed and cycling in CV but not GF mice (Fig. 2A). HDACs repress transcription by deacetylating histones but can also target non-histone proteins. We examined expression of multiple *Hdac* genes (*Hdac1*-*Hdac11*), finding that *Hdac1* and *Hdac3* were more highly expressed in IECs than the other *Hdac* genes (fig. S3A). However, only *Hdac3* showed differential expression in CV and GF mice, with lowered abundance of *Hdac3* transcripts and HDAC3 protein in IECs from GF as compared to CV mice (Fig. 2B,C; fig. S3B,C). *Hdac3* expression increased in IECs after conventionalization of adult GF mice (Fig. 2D), and was selectively triggered by monocolonization of GF mice with the commensal bacterial species *Bacteroides thetaiotaomicron* (fig. S3D). Elevated *Hdac3* expression in CV mice required the Toll-like receptor adaptor protein MyD88 (*14*) (fig. S3E), and accordingly, *MyD88*^*-/-*^ mice showed dampened rhythms in small intestinal histone acetylation (fig. S3F).

**Figure 2:**
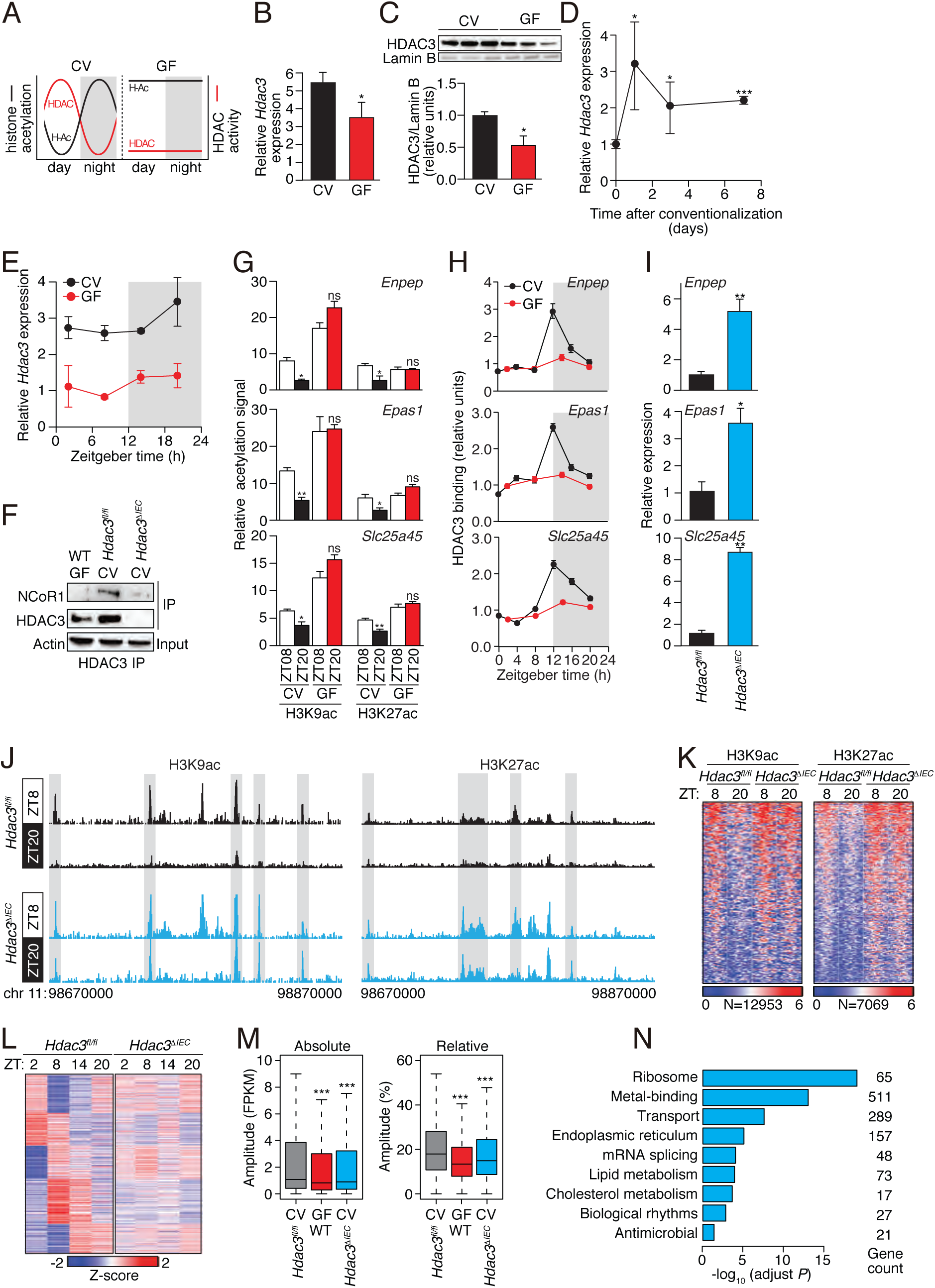
The intestinal microbiota programs the diurnal rhythmicity of epithelial histone acetylation and gene expression through HDAC3. **(A)** The diurnal histone acetylation pattern in CV and GF mice suggests a loss of HDAC function in GF mice. **(B)** RT-qPCR analysis of *Hdac3* transcripts in IECs from CV and GF mice. N=3 mice per group. **(C)** Western blot of HDAC3 and Lamin B (loading control) in IECs from protein in intestinal epithelial cells from CV and GF mice. Bands were quantified by scanning densitometry and normalized to the Lamin B band intensity. **(D)** RT-qPCR analysis of *Hdac3* transcript abundance in epithelial cells from GF mice and GF mice following conventionalization. N=3 mice per group. **(E)** RT-qPCR analysis of *Hdac3* transcripts in IECs from CV and GF mice across a 24-hour cycle. N=3 mice per group. **(F)** Co-immunoprecipitation of endogenous HDAC3 and NCoR1 in IECs from GF WT mice and CV *Hdac3*^*fl/fl*^ and *Hdac3*^*ΔIEC*^ mice. N=3 pooled biological replicates per lane. **(G**,**H)** Relative abundance of H3K9ac and H3K27ac marks **(G)** and bound HDAC3 **(H)** at the promoters of *Enpep, Epas1* and *Slc25a45* in CV and GF as determined by ChIP-qPCR analysis. N=3 mice per time point per group. **(I)** RT-qPCR analysis of *Enpep, Epas1* and *Slc25a45* transcript abundance in IECs from *Hdac3*^*fl/fl*^ and *Hdac3*^*ΔIEC*^ mice at ZT8. N=3 mice per group. (**J)** Genome browser view of the 200 kb region surrounding the *Nr1d1* locus, showing ChIP-seq analysis of H3K9ac and H3K27ac marks in small intestinal epithelial cells from *Hdac3*^*fl/fl*^ and *Hdac3*^*ΔIEC*^ mice. N=3 pooled biological replicates per library. **(K)** Heat maps of H3K9ac and H3K27ac signals (log (reads at 50 bp windows)) from −1 to +1 kb surrounding the centers of all cycling H3K9ac and H3K27ac peaks. **(L)** Heat map showing diurnal gene expression patterns in IECs from *Hdac3*^*fl/fl*^ and *Hdac3*^*ΔIEC*^ mice across a 24-hour cycle by RNA-seq. N=3 pooled biological replicates per library. **(M)** Absolute (Fragments Per Kilobase of transcript per Million mapped reads, FPKM) or relative oscillating amplitudes of transcripts in GF *WT* mice and CV *Hdac3*^*fl/fl*^ and *Hdac3*^*ΔIEC*^ mice. N=3 mice per group. **(N)** Enriched GO categories of genes with decreased cycling amplitudes as determined by DAVID. \**P*<0.05; \*\**P*<0.01; ***P<0.001; ns, not significant by two-tailed *t*-test. Means±SEM (error bars) are plotted. GF, germ-free; CV, conventional; ZT, Zeitgeber time.

We reasoned that if HDAC3 is essential for microbiota-driven diurnal rhythms in epithelial histone acetylation, then HDAC3 expression, genome recruitment, or enzymatic activity should be diurnally rhythmic. *Hdac3* transcript abundances showed only a modest diurnal oscillation (Fig. 2E), suggesting that the rhythmicity in IEC histone acetylation was not driven by rhythmic *Hdac3* expression. In liver and adipose tissue, rhythmicity in histone acetylation is generated when HDAC3 complexes with the nuclear receptor co-repressor (NCoR) (*12, 15-18*), which is recruited to chromatin in a diurnally rhythmic manner by components of the circadian clock (*12*). We therefore postulated that rhythms in IEC chromatin acetylation might be generated by rhythmic chromatin recruitment of HDAC3 through binding to NCoR.

In support of this idea, we found that HDAC3 complexed with NCoR in IECs, and that binding was markedly reduced in IECs from GF mice (Fig. 2F). Thus, the microbiota facilitates formation of the HDAC3-NCoR complex in IECs. We next assessed whether HDAC3 was rhythmically recruited to target genes by studying three genes, *Enpep, Epas1*, and *Slc25a45*, that showed pronounced microbiota-dependent rhythms in H3K9ac and H3K27ac marks (Fig. 2G; fig. S4). As predicted, HDAC3 was recruited to each of these genes in a diurnally rhythmic manner that depended on the microbiota (Fig. 2H). Expression of each gene was higher in mice with an IEC-specific deletion of *Hdac3 (Hdac3*^Δ*IEC*^)(Fig. 2I), consistent with the canonical function of HDAC3 as a co-repressor through histone deacetylation (*18*). Thus, the microbiota is required for diurnally rhythmic recruitment of HDAC3 to target gene promoters. Together, these data argue that microbiota-driven formation of the HDAC3-NCoR complex promotes the diurnally rhythmic recruitment of HDAC3 to target genes and consequent rhythmic histone deacetylation in CV IECs.

To determine whether HDAC3 drives genome-wide rhythmic histone acetylation in IECs, we detected H3K9ac and H3K27ac marks by ChIP-seq in IECs from *Hdac3*^Δ*IEC*^ mice. Both marks oscillated diurnally in *Hdac3*^*fl/fl*^ IECs but showed dampened oscillations in the *Hdac3*^Δ*IEC*^ IECs (Fig. 2J,K). We also performed RNA-seq of *Hdac3*^*fl/fl*^ and *Hdac3*^Δ*IEC*^ IECs across a circadian cycle. Rhythmic oscillations in the abundances of 2729 transcripts were dampened in *Hdac3*^Δ*IEC*^ IECs, approximating the amplitudes seen in GF mice (Fig. 2L,M). Genes having dampened oscillations in *Hdac3*^Δ*IEC*^ mice were enriched for nutrient transport and lipid metabolism pathways (Fig. 2N), similar to the enrichment of these pathways among genes with oscillating acetylation marks (Fig. 1E). Rhythmic expression of the clock genes *Bmal1* (*Arnt1*), *Per2*, and *Nr1d1* (*Rev-erba*) was maintained in *Hdac3*^Δ*IEC*^ IECs (Fig. 3A,B), indicating that the core clock mechanism remains intact. Thus, the intestinal microbiota regulates genome-wide diurnal rhythms in IEC acetylation and gene expression through HDAC3.

**Figure 3:**
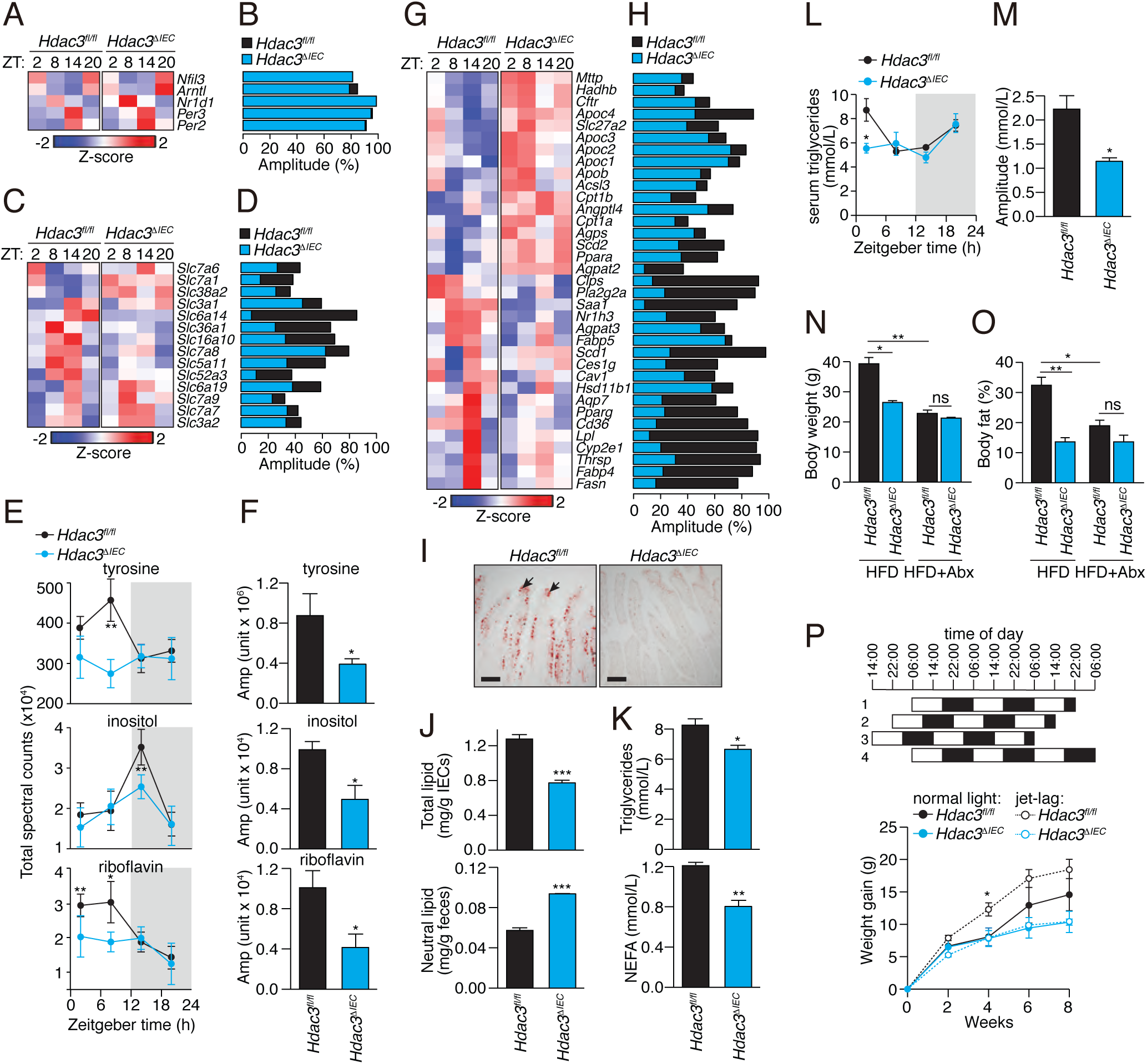
Intestinal epithelial HDAC3 regulates the diurnal rhythmicity of nutrient uptake and controls lipid absorption in the intestine. **(A)** Diurnal expression pattern of circadian clock genes in *Hdac3*^*fl/fl*^ and *Hdac3*^*ΔIEC*^ mice IECs are represented by a heat map, with amplitudes shown in **(B). (C)** Diurnal expression patterns of transporter genes in *Hdac3*^*fl/fl*^ and *Hdac3*^*ΔIEC*^ mice IECs are represented by a heat map, with amplitudes shown in **(D). (E)** Diurnal oscillations of serum tyrosine, inositol and riboflavin in *Hdac3*^*fl/fl*^ and *Hdac3*^*ΔIEC*^ mice with the amplitudes shown in **(F)**. N=3 mice per group. \**P*<0.05 by two-tailed *t*-test. **(G)** Diurnal expression pattern of lipid metabolic genes in IECs from *Hdac3*^*fl/fl*^ and *Hdac3*^*ΔIEC*^ mice IECs is represented by a heat map, with amplitudes shown in **(H). (I)** Oil red O staining of lipids in small intestines of *Hdac3*^*fl/fl*^ and *Hdac3*^*ΔIEC*^ mice fed a HFD. Black arrows indicate epithelial cells. Scale bars=100 µm. **(J)** Quantification of total lipids in IECs and neutral lipids in feces from *Hdac3*^*fl/fl*^ and *Hdac3*^*ΔIEC*^ mice fed a HFD. N=6, 5 mice per group. \*\*\**P*<0.001 by two-tailed *t*-test. **(K)** Quantification of triglycerides and free fatty acids in serum from *Hdac3*^*fl/fl*^ and *Hdac3*^*ΔIEC*^ mice fed a HFD. N=6, 5 mice per group. \**P*<0.05; \*\**P*<0.01 by one-tailed *t*-test. **(L)** Diurnal oscillations of serum triglyceride concentrations in *Hdac3*^*fl/fl*^ and *Hdac3*^*ΔIEC*^ mice fed a chow diet, with ampitudes shown in **(M)**. N=3 mice per group. \**P*<0.05 by one-tailed *t*-test. **(N**,**O)** Weight **(N)** and body fat percentage **(O)** of *Hdac3*^*fl/fl*^ and *Hdac3*^*ΔIEC*^ mice fed a HFD with or without antibiotics for 10 weeks. N=5 mice per group. \**P*<0.05; \*\**P*<0.01; ns, not significant by one-tailed *t*-test**. (P)** Weight gain in mice fed a HFD with or without jet lag. Jet lag was induced by an 8 hour light cycle shift every three days. N=5 mice per group). \**P*<0.05 two-tailed *t*-test. Means±SEM (error bars) are plotted. ZT, Zeitgeber time; HFD, high fat diet.

The RNAseq analysis identified several nutrient transporter genes with diurnally rhythmic expression that was dampened in *Hdac3*^Δ*IEC*^ mice (Fig. 3C,D). These genes included *Slc5a11, Slc52a3*, and *Slc16a10*, which transport aromatic amino acids, inositol, and riboflavin respectively (*19-22*). Accordingly, *Hdac3*^Δ*IEC*^ mice showed dampened daily rhythms in the serum concentrations of several metabolites, including tyrosine (an aromatic amino acid), inositol, and riboflavin (Fig. 3E,F; fig. S5). This suggests that epithelial HDAC3 regulates nutrient uptake in the small intestine.

Genes with functions in lipid metabolism also showed dampened diurnal expression rhythms in the *Hdac3*^Δ*IEC*^ mice. Seventeen of these genes showed increased overall expression in the *Hdac3*^Δ*IEC*^ mice, including *Acsl3, Cpt1a/b, Ppara*, and *Agpat2*, which participate in fatty acid β-oxidation and lipid biosynthesis (Fig. 3G,H). In contrast, eighteen genes that regulate lipid uptake and processing had decreased overall expression in the *Hdac3*^Δ*IEC*^ mice. These genes included *Cd36*, encoding a fatty acid transporter (*23*), *Fabp4/5*, encoding fatty acid binding proteins (*24*), and *Scd1*, encoding a stearoyl-coenzyme A-desaturase (*25*)(Fig. 3G,H). Consistent with the reduced *Cd36* expression, *Hdac3*^Δ*IEC*^ mice had lowered IEC total lipid concentrations and increased fecal lipid concentrations (Fig. 3I,J). Our finding of decreased lipid uptake differs from a prior report of increased lipid stores in IECs of *Hdac3*^Δ*IEC*^ mice (*26*). However, we note that the mice in the prior study were fed a defined safflower lipid diet that differs from the HFD used in our studies (TestDiet AIN-76A), possibly explaining the discordant findings. We also observed lower concentrations of serum triglycerides and non-esterified fatty acids (Fig. 3K) and dampened diurnal rhythms in serum triglyceride concentrations in the *Hdac3*^Δ*IEC*^ mice (Fig. 3L,M). Thus, HDAC3 enhances expression of the lipid transporter CD36, stimulates lipid uptake by IECs, and promotes diurnal rhythmicity in serum triglyceride concentrations.

The lowered lipid uptake into *Hdac3*^Δ*IEC*^ IECs suggested that the *Hdac3*^Δ*IEC*^ mice might be protected against high fat diet-induced obesity. Indeed, when placed on a high-fat, Western-style diet (HFD) for 10 weeks, the *Hdac3*^Δ*IEC*^ mice maintained lower body weights and had lower body-fat percentages than *Hdac3*^*fl/fl*^ mice (Fig. 3N,O; fig. S6A), which accords with a prior report (*26*). The *Hdac3*^Δ*IEC*^ mice also had increased glucose tolerance, decreased insulin resistance, smaller epididymal fat pads, and less liver fat accumulation (fig. S6B-D). The metabolic differences between *Hdac3*^*fl/fl*^ and *Hdac3*^Δ*IEC*^ mice were not due to altered physical activity, energy utilization (fig. S7A), food intake, (fig. S7B), or altered microbiota taxonomic composition (fig. S8), which were similar between the two groups. Thus, epithelial HDAC3 promotes HFD-induced obesity in mice, likely by enhancing intestinal lipid absorption.

The microbiota is an essential driver of diet-induced obesity, and thus GF mice have lower body fat percentages and are protected from HFD-induced obesity relative to CV mice (*2*). The lowered expression and dampened rhythmic recruitment of HDAC3 to epithelial chromatin in GF mice suggested a causal role for epithelial HDAC3 in microbiota-dependent obesity. Depletion of the microbiota through antibiotic treatment produced lowered body weight and body fat percentages in the *Hdac3*^*fl/fl*^ mice, resulting in body compositions that did not differ significantly from those of *Hdac3*^Δ*IEC*^ mice (Fig. 3N,O). Thus, epithelial HDAC3 is necessary for the microbiota to promote obesity in mice.

Chronic circadian light cycle disruptions exacerbate HFD-induced obesity in mice (*8*). Consistent with prior findings (*8*), *Hdac3*^*fl/fl*^ mice subjected to experimental jet lag (an 8-hour shift every 3 days) gained weight faster than *Hdac3*^*fl/fl*^ mice on a normal day-night light cycle (Fig. 3P). However, *Hdac3*^Δ*IEC*^ mice were protected from the metabolic effects of jet lag and gained weight at a similar rate as *Hdac3*^Δ*IEC*^ mice on a normal light cycle. Their overall weight gain was also lower than *Hdac3*^*fl/fl*^ mice that were either jet-lagged or under a normal light cycle. Thus, epithelial HDAC3 enhances jet lag-induced obesity in mice.

We next investigated the molecular mechanisms by which HDAC3 regulates the rhythmic expression of lipid metabolic pathways in the small intestinal epithelium. Canonically, HDAC3 acts as a corepressor through its histone deacetylation activity. The elevated expression of genes involved in fatty acid β-oxidation and lipid biosynthesis in *Hdac3*^Δ*IEC*^ mice (Fig. 3G) suggested that HDAC3 functions canonically as a corepressor of these genes. Accordingly, H3K9ac marks at *Cpt1a, Acsl3, Ppara*, and *Agpat2* oscillated diurnally in conventional mice but were constitutively elevated and non-rhythmic in GF mice and conventional *Hdac3*^Δ*IEC*^ mice(Fig. 4A,B; fig. S9A,B), consistent with HDAC3 promoting histone deacetylation that suppresses transcription of these genes.

**Figure 4:**
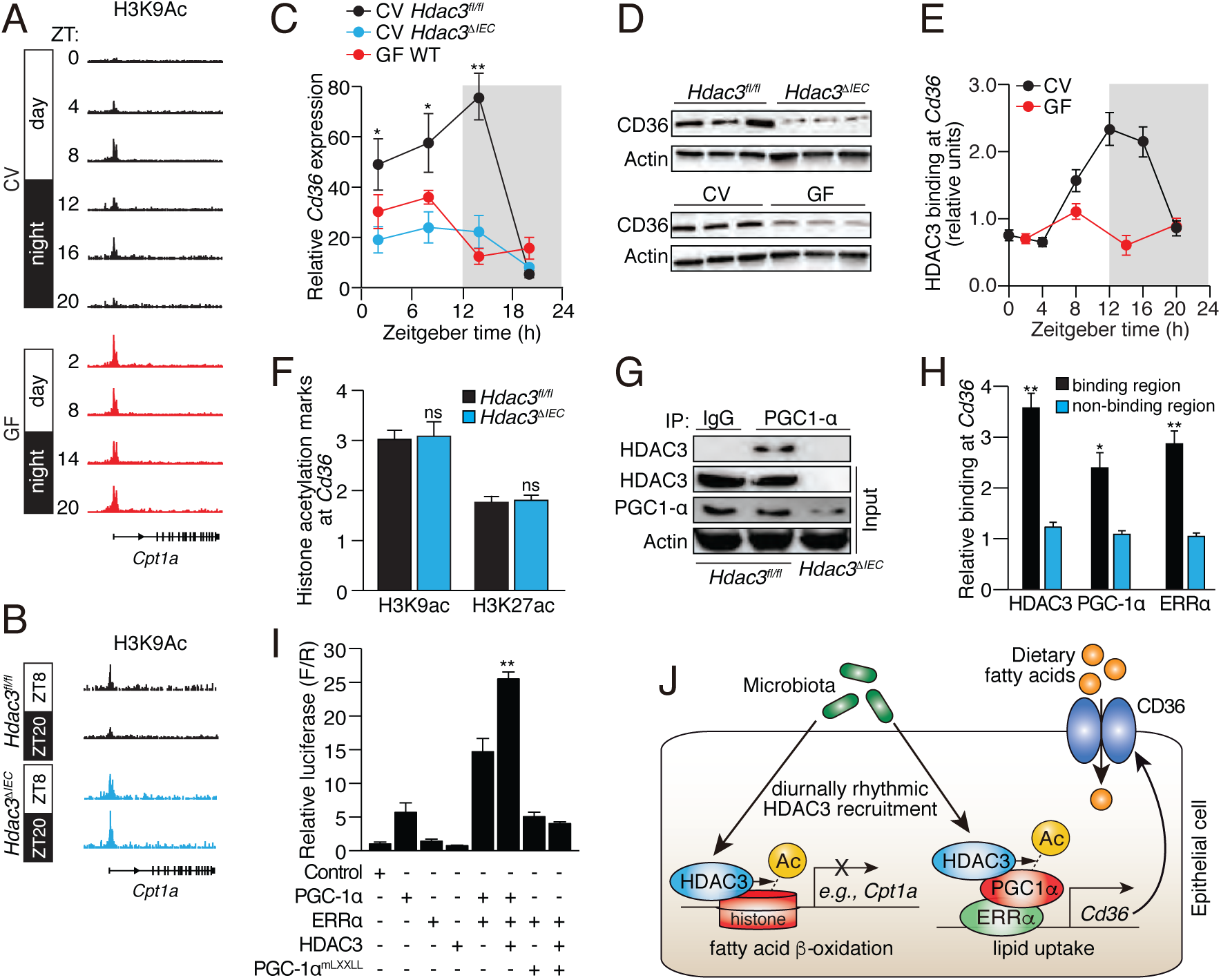
Epithelial HDAC3 regulates expression of the lipid transporter gene *Cd36* by coactivating ERRα through PGC-1α. **(A,B)** Genome browser view of the *Cpt1a* locus, showing H3K9ac marks in small intestinal epithelial cells from CV and GF WT mice **(A)**, and CV *Hdac3*^*fl/fl*^ and *Hdac3*^*ΔIEC*^ mice **(B). (C,D)** Diurnal expression of *Cd36* transcript **(C)** and protein **(D)** in IECs from *Hdac3*^*fl/fl*^ CV mice, *Hdac3*^*ΔIEC*^ CV mice and GF WT mice. N=3 mice per group. **(E)** ChIP-qPCR analysis of HDAC3 binding at the *Cd36* promoter in CV and GF mice across a day-night cycle. **(F)** ChIP-qPCR analysis of H3K9ac and H3K27ac marks at the *Cd36* promoter in IECs from *Hdac3*^*fl/fl*^ and *Hdac3*^*ΔIEC*^ mice. **(G)** Co-immunoprecipitation of endogenous HDAC3 and PGC-1α in IECs from *Hdac3*^*fl/fl*^ and *Hdac3*^*ΔIEC*^ mice. N=3 pooled biological replicates per lane. **(H)** ChIP-qPCR analysis of HDAC3, PGC-1α and ERRα shows co-localization at the *Cd36* promoter in IECs from CV WT mice. “Binding region” refers to a known binding site for each protein at the *Cd36* locus in adipose tissue (*17*). N=3 mice per group. **(I)** Luciferase reporter assay of transcription driven by a *Cd36* promoter, demonstrating combinatorial effects of HDAC3, PGC-1α and ERRα. Empty vector was used in the control group. N=3 mice per group. **(J)** Model showing how the intestinal microbiota regulates diurnal rhythms in epithelial metabolic pathways through HDAC3. \**P*<0.05; \*\**P*<0.01; ns, not significant by two-tailed *t*-test. Means±SEM (error bars) are plotted. ZT, Zeitgeber time; CV, conventional; GF, germ-free.

By contrast, genes that control lipid uptake, transport, and processing, including *Cd36, Scd1*, and *Fabp4*, showed lowered expression in *Hdac3*^Δ*IEC*^ IECs (Fig. 3G). This suggested that HDAC3 regulates expression of these genes either indirectly or through a non-canonical coactivation mechanism. Several of the genes in this group (including *Cd36, Scd1*, and *Fabp4*) are induced by the microbiota through the circadian transcription factor NFIL3 (*7*), suggesting that HDAC3 might regulate their expression by controlling *Nfil3* expression. However, *Nfil3* expression was not markedly altered in *Hdac3*^Δ*IEC*^ mice and remained rhythmic (Fig. 3A,B), arguing against a role for NFIL3.

To further understand the mechanism of gene activation by HDAC3 we studied the fatty acid transporter gene *Cd36*. *Cd36* expression in the intestinal epithelium oscillated diurnally in conventional *Hdac3*^*fl/fl*^ mice, but the oscillations were dampened and overall transcript and protein levels were lowered in germ-free mice and conventional *Hdac3*^Δ*IEC*^ mice (Fig. 4C,D). This finding supports the idea that the microbiota promotes *Cd36* expression and hence lipid absorption through HDAC3. HDAC3 was recruited rhythmically to *Cd36* in a microbiota-dependent manner (Fig. 4E), but H3K9ac and H3K27ac marks at *Cd36* were not altered in *Hdac3*^Δ*IEC*^ mice (Fig. 4F). These data suggest that HDAC3 regulates *Cd36* expression directly but not through histone acetylation.

In adipose tissue, HDAC3 functions as a co-activator of estrogen-related receptor α (ERRα)(*17*). Transcriptional activation by ERRα requires the proliferator-activated receptor gamma coactivator 1 α (PGC1α), which is in turn activated by HDAC3-mediated deacetylation (*17, 27*). We therefore considered whether a similar mechanism might underlie HDAC3 activation of *Cd36* expression in IECs. Supporting this idea, analysis of published genome-wide ChIP-seq data from adipose tissue showed simultaneous recruitment of HDAC3 and ERRα to multiple *Cd36* promoter and enhancer sites ((*17*); Fig. S10A). Further, HDAC3 co-immunoprecipitated with PGC1α in IECs, indicating a direct physical interaction between the proteins in epithelial cells (Fig. 4G). HDAC3, PGC-1α and ERRα colocalized at *Cd36* in IECs (Fig. 4H), and HDAC3 markedly increased the transcriptional activity of ERRα at *Cd36* (Fig. 4I; Fig. S10B). This increase occurred with wild-type PGC1α, but not with a mutant PGC1α that cannot interact with ERRα (*17*). Thus, the microbiota promotes diurnally rhythmic expression of intestinal *Cd36* through HDAC3 and its function as a co-activator of ERRα (Fig. 4J).

Circadian rhythmicity is a defining characteristic of mammalian metabolism that coordinates expression of cellular metabolic machinery with environmental light cycles. Our finding that the intestinal microbiota programs the daily rhythmic expression of small intestinal metabolic networks illuminates an essential role for the microbiota in regulating host metabolism, and indicates that the microbiome, the circadian clock, and the mammalian metabolic system are tightly co-evolved. We identify epithelial HDAC3 as a key mechanism that integrates inputs from the microbiota and circadian light cycles and conducts these signals to host metabolic networks. The microbial-circadian interaction mediated by HDAC3 regulates intestinal lipid uptake, and is therefore essential for the development of diet-induced obesity and the worsening of obesity by jet lag. Our results suggest how disruption of microbiota-clock interactions, such as through antibiotic treatment or chronic circadian disruptions including jet lag, could promote human metabolic disease. These findings also point to new avenues for treating metabolic disease by chemical or microbiological targeting of HDAC3.

## Supporting information

Kuang et al Supplemental Information

## Acknowledgements

We thank B. Hassell and M. Robinson for assistance with mouse experiments. This work was supported by NIH grant R01 DK070855 (L.V.H.), Welch Foundation Grant I-1762 (L.V.H.), the Helen D. Bader Center for Research on Arthritis and Autoimmune Diseases (L.V.H.), and the Howard Hughes Medical Institute (L.V.H.). Z.K. was supported by NIH T32 AI005284. All data and code to understand and assess the conclusions of this research are available in the main text, supplementary materials, and via the Gene Expression Omnibus repository with accession number XXXX.

