## Supplementary material for "The intestinal microbiota programs diurnal rhythms in host metabolism through histone deacetylase 3": Kuang et al Supplemental Information

### Supporting Online Material

### Materials and Methods

#### Mice

Wild-type C57BL/6, *Hdac3*$^{fl/fl}$ (*28*), *Hdac3*$^{\Delta IEC}$, and *Myd88*$^{-/-}$ mice were bred and maintained in the SPF barrier at the University of Texas Southwestern Medical Center. *Hdac3*$^{fl/fl}$ mice (*28*) were used to generate intestinal epithelial cell (IEC)-specific *Hdac3* knockout mice (*Hdac3*$^{\Delta IEC}$) by crossing with a mouse expressing Cre recombinase under the control of the IEC-specific *Villin* promoter (*29*). Germ-free (GF) C57BL/6 mice were bred and maintained in the gnotobiotic mouse facility at the University of Texas Southwestern Medical Center as described (*30*). All mice were housed under a 12-hour light and 12-hour dark cycle unless otherwise specified. Mice were fed *ad libitum*. All experiments were performed using protocols approved by the Institutional Animal Care and Use Committee of the UT Southwestern Medical Center.

#### Jet lag experiments

Eight week old *Hdac3*$^{fl/fl}$ and *Hdac3*$^{\Delta IEC}$ mice were housed in ventilated, light-tight cabinets on a 12-hour light and 12-hour dark cycle (Phenome Technologies). After acclimation for 3 days, light cycles were changed for mice subjected to experimental jet lag while the light cycles of control groups were left unchanged. Every 3 days, lights were turned on 8 hours earlier than the previous setting while maintaining a 24-hour light/dark cycle thereafter.

**Diet**

Mice were fed a regular chow diet (LabDiet 5KA1) containing 22% protein, 16% fat and 62% carbohydrates, or a Western style high fat diet (HFD) (TestDiet AIN-76A) containing 16% protein, 40% fat and 44% carbohydrates.

**Metabolic studies**

Mice were fed the HFD for 10 weeks before metabolic analysis. Antibiotic-treated mice were given an antibiotic cocktail containing 1 g/L streptomycin, 1 g/L gentamycin, 1 g/L neomycin, 1 g/L metronidazole and 0.5 g/L vancomycin throughout the HFD treatment. Body composition was measured with an EchoMRI™-100H analyzer. Blood glucose was measured using a One Touch Ultra2 glucose meter. Serum triglycerides were quantified using Infinity Triglycerides Liquid Stable Reagent (Thermo Scientific TR22421). Free fatty acids were measured using the Wako NEFA-HR(2) reagent (Wako 434-91795, 436-91995, 270-77000). Circadian oscillations of serum triglycerides were measured in mice fed *ad libitum* on a chow diet. Steady-state serum triglycerides, free fatty acids, and glucose tolerance were measured after an overnight fast. The insulin tolerance test was performed after a 4 hour fast. Mice were injected intraperitoneally with 2 mg/g body weight of D-(+)-glucose (Sigma G8769), or 0.5 U/kg body weight insulin (Eli Lilly, Humulin R) and blood glucose was measured at time 0, 15, 30, 60 and 100 minutes after injection. Real-time metabolic cage analysis was performed with the TSE Labmaster System.

**Oil Red O detection of cellular lipids**

Small intestines from $Hdac3^{fl/fl}$ and $Hdac3^{\Delta IEC}$ mice fed a HFD were rinsed with PBS and fixed in 10% formalin at room temperature for 2 hours. Tissues were rinsed with 2% sucrose in PBS and snap frozen in optimum cutting temperature (OCT) compound (Fisher 23-730.571). 7 µm sections were cut and mounted to Superfrost Plus Micro Slides (VWR). Oil Red O stock solution (0.5 g/100 mL dissolved in isopropanol) was diluted with distilled water (6:4, V/V) and filtered through a 0.2 µm filter to make the working Oil Red O solution. Tissue sections were post-fixed with 10% formalin for 15 min, dipped in 60% isopropanol, and stained with the Oil Red O working solution for 30 min. Slides were then destained in 60% isopropanol and rinsed under running tap for 2 min. Images were captured using a Zeiss AxioImager M1 microscope.

**Lipid quantification in intestinal epithelial cells**

Ileal IECs from *Hdac3*$^{fl/fl}$ and *Hdac3*$^{\Delta IEC}$ mice on HFD were isolated using 10 mM EDTA as previously described (*7*). 10-20 mg IECs from each sample were used for lipid extraction using the Lipid Extraction Kit (Cell Biolabs STA-162). Extracted lipids were air dried, resuspended in 100 µl cyclohexane, and quantified using a Lipid Quantification Kit (Cell Biolabs STA-613). Lipid concentrations were normalized to cell weight.

**Neutral lipid quantification in fecal pellets**

Fresh fecal pellets from *Hdac3*$^{fl/fl}$ and *Hdac3*$^{\Delta IEC}$ mice fed the HFD were collected and weighed. Fecal lipids were extracted using a Lipid Extraction Kit (Cell Biolabs STA-612). Extracted lipids were air dried, resuspended in 100 µl isopropanol, and quantified using a Lipid Quantification Kit for neutral lipids (Cell Biolabs STA-617). Concentrations were normalized to feces weight.

**Liquid chromatography (LC) with mass spectrometry (MS) analysis**

Freshly prepared serum was quenched immediately with the same volume of chilled (-20°C) HPLC-grade methanol and spun for 20 min at 16,000 RCF in a tabletop centrifuge. The supernatants were collected and dried in a vacuum concentrator. Extracted metabolites were resuspended in detection buffers, separated chromatographically on a $C_{18}$ column, and detected using an AB SCIEX 3200 QTRAP triple quadrupole mass spectrometer with different targeted LC-MS/MS methods as previously established (*31, 32*).

**Laser capture microdissection and RNA purification**

Laser capture microdissection was performed as previously described (*33*). Briefly, a 5 cm length of mouse ileum was washed and snap-frozen in OCT compound (Fisher 23-730.571). 7 μm frozen sections were cut and fixed in 70% ethanol and sequentially stained with Methyl Green and eosin. Laser capture microdissection of IECs was performed using an Arcturus PixCell IIe system. 5000-10000 cells were obtained from each section in one hour, and RNA was immediately extracted from the captured IECs by incubating with 14 μl RNA extraction buffer from the PicoPure RNA Isolation Kit (Life Technology 12204-01) for 30 min. Extracted RNA was purified using PicoPure RNA Isolation Kit.

**Quantitative real-time PCR**

cDNA was synthesized from purified RNA using M-MLV Reverse Transcriptase (Fisher 28025021), qRT-PCR was performed using Platinum SYBR Green qPCR SuperMix (Fisher 11733046) on a QuantStudio 7 Flex Real-Time PCR System (Applied Biosystems). Expression levels were calculated relative to the abundance of *Gapdh* transcripts. Primer sequences are listed in Table S1.

**Chromatin immunoprecipitation (ChIP)**

Mouse ileum was washed with ice-cold PBS and IECs were extracted in 10 mM EDTA. The ChIP assay was carried out as previously described (*34, 35*) with a few modifications. Briefly, cells were washed with ice-cold PBS twice and fixed in 1% formaldehyde at room temperature for 10 min. Cells were then quenched in 125 mM glycine for 10 min, washed twice with PBS, resuspended in 0.5 ml lysis buffer (20 mM Tris-HCl, pH 8, 60 mM KCl, 1 mM EDTA, 0.5% NP-40 with protease inhibitors), and incubated at 4°C for 15 min. Nuclei were pelleted and resuspended in 250 μl RIPA buffer (Thermo Scientific 89900) with protease inhibitors and sonicated with a Bioruptor Pico sonication device (Diagenode) for 20 cycles (30 sec on, 30 sec off). The supernatant was pre-cleared with 10 μl protein G magnetic beads for 1 h and then incubated with 3 μg of primary antibodies overnight at 4°C. The antibodies used were: anti-H3K9ac (Abcam ab4441), anti-H3K27ac (Abcam ab4729), anti-HDAC3 (Abcam ab7030), anti-PGC-1α (Millipore, ST1204), anti-ERRα (Millipore 17-603). 50 μl protein G beads were added and incubated for 1.5 h at 4°C. Beads were washed four times with LiCl wash buffer (100 mM Tris-HCl pH 7.5, 500 mM LiCl, 0.5% NP-40, 0.5% sodium deoxycholate) and finally with TE buffer. DNA was eluted in 150 μl TES buffer (TE with 1% SDS, 150 mM NaCl, and 5 mM dithiothreitol) by resuspending the beads at 65°C for 8 h. DNA was purified using ChIP DNA Clean & Concentrator (Zymo Research D5205). ChIP DNA was either used for sequencing or quantative PCR (qPCR). For ChIP-qPCR, relative enrichment was calculated by first normalizing all the signals from immunoprecipitated DNA to the signals from input DNA and further normalizing signals from binding regions to control regions. Binding region refers to known binding sites for each mark at the gene of interest and control region refers to the neighboring regions that lack binding sites.

**Western blot**

Mouse ileum tissues were washed with ice-cold PBS and IECs were isolated in 10 mM EDTA as previously described (*7*). Cells were washed twice with PBS and resuspended in 250 μl RIPA buffer (Thermo Scientific 89900) with protease inhibitors and sonicated with a Bioruptor Pico sonication device (Diagenode) for 10 cycles (30 sec on, 30 sec off). Supernatant protein concentrations were measured using the Pierce BCA Protein Assay Kit (Thermo 23225) and normalized across samples. Lysates were separated on 4-20% gradient SDS-PAGE gels and transferred to PVDF membranes. Membranes were blocked with 5% nonfat milk in TBS-T buffer (0.1% Tween-20 in Tris-buffered saline) for 1 h at room temperature then sequentially incubated with primary antibodies: anti-CD36 (ThermoFisher PA1-16813), anti-actin (Sigma, A5060), anti-Lamin B (abcam ab133741), anti-PGC-1α (Millipore, ST1202), anti-HDAC3 (Abcam ab7030), and appropriate HRP-conjugated secondary antibodies. Protein bands were visualized using a Bio-Rad ChemiDoc system.

**Co-immunoprecipitation**

Mouse ileum tissues were washed with ice-cold PBS and IECs were isolated in 10 mM EDTA as previously described (*7*). Cells were washed twice with PBS and resuspended in NP40 IP lysis buffer (20 mM Tris-HCl, pH 8.0, 150 mM NaCl, 10% glycerol, 2 mM EDTA, 0.1 % NP40, 10 mM NaF, 1 mM DTT, 2 mM phenylmethylsulfonylfluoride and protease inhibitors). Lysates were sonicated for 10 cycles (30 sec on, 30 sec off) and supernatants were pre-cleared with protein G beads. After normalizing protein concentration, 3 μg of primary antibodies (anti-PGC-1α (Millipore, ST1204); anti-HDAC3 (Abcam ab7030); anti-rabbit IgG (ThermoFisher 02-6102)) were added and incubated at 4°C for 2 hours. Next, protein G magnetic beads were added and incubated at 4°C for 1 hour. Beads were washed with NP40 IP lysis buffer four times and samples were eluted in 2X SDS loading buffer at 95°C for 5 min. Eluted proteins were separated on a 4-20% gradient SDS-PAGE gel and transferred to a PVDF membrane. The membrane was incubated with primary antibodies overnight and detected using a Bio-Rad ChemiDoc system. Input proteins were detected with anti-HDAC3 (Abcam, ab187945), anti-PGC-1α (Millipore, ST1202), anti-NCoR1 (Abcam, ab3482), and anti-actin antibody (Sigma, A5060).

**Immunostaining**

Mouse ileum was washed with PBS, fixed in Bouin's fixative overnight at 4°C and embedded in paraffin. Sections were washed in xylene twice and rehydrated in decreasing concentrations of ethanol (100%, 95%, 70%, 50%, 0%). Slides were boiled in 10 mM sodium citrate for 15 min and washed in PBS twice. Slides were blocked with 10% FBS, 1% BSA, 1% Trixon X-100 in PBS for 1 hour and incubated with primary antibody (anti-HDAC3, ab7030, 1:200 dilution) at 4°C overnight. Cy3 anti-rabbit secondary antibody (Fisher 711-165-152) was diluted 1:400 and applied to slides for 1 hour at room temperature in the dark. Slides were washed and mounted with DAPI Fluoromount-G (Southern Biotechnology 0100-20). Images were captured using a Zeiss AxioImage MI microscope.

**Conventionalization and monoassociation of GF mice**

For conventionalization, ~50 mg feces was collected from a conventional wild-type mouse and suspended in 1 ml PBS. Fecal debris was pelleted and 200 µl of the supernatant was used for oral gavage of each GF mouse. Mice were sacrificed 1, 3 and 7 days after colonization. For monoassociations, GF mice were colonized with 2 x $10^9$ colony forming units (cfu) of log phase *Bacteroides thetaiotaomicron* (VPI-5482), *Enterococcus faecalis* (ATCC-29212), *Escherichia coli* K235 (ATCC-13027) or *E. coli* O127:K63 (ATCC-12740), or with 1 x $10^9$ cfu of log phase *Salmonella typhimurium* (strain 1433) through oral gavage. Mice were sacrificed 3 days after colonization.

**Luciferase assay**

A 1296 bp fragment of *Cd36* promoter (variant 5, NM_001159556.1) was fused to a firefly luciferase reporter (pGL3 promoter vector). ~1.5 kb fragments of *Cd36* enhancers were fused to pGL4.24 enhancer luciferase vector. 2 x $10^4$ MODE-K cells were seeded into each well of a 96-well plate one day before transfection. 90 ng of empty vector, or protein expression vectors were transfected together with 30 ng *Cd36* reporter vector and 10 ng Renilla reporter vector. 24 hours after transfection, luciferase activities were determined using the Dual-Glo Luciferase Assay System (Promega E2920) following the manufacturer's protocol. The Renilla luciferase co-reporter was used to normalize luciferase activity.

**RNA-seq and data analysis**

RNA was extracted, pooled and purified from ileal epithelial cells of three mice at each time point across a 24-hour circadian cycle. RNA quality was examined by an Agilent 2100

Bioanalyzer. Libraries were prepared using a TruSeq RNA sample preparation kit v2 (Illumina) and sequencing was performed on Illumina NextSeq for single-end 75 bp length reads. Data were analyzed as previously described (*7*). Sequence data were mapped against the mm10 genome using TopHat (*36*) and FPKMs were generated using Cuffdiff (*36*) with default parameters. Circadian oscillation was analyzed by JTK (*37*). Absolute amplitudes were directly output by JTK and relative amplitudes were calculated by dividing absolute amplitudes to the average expression levels. Expression levels of genes with circadian adjusted $P$ Values < 0.05 (by JTK) or log2 fold changes of circadian amplitudes > 1 or log2 fold changes of FPKMs > 1 or < -1 were plotted in heat maps. Gene Ontology analysis was performed using the online tool DAVID Bioinformatics Resources (https://david.ncifcrf.gov/).

**ChIP-seq and data analysis**

ChIP-seq libraries were generated using a KAPA Hyper Prep Kit (kapabiosystems KK8502) and library sizes and concentrations were evaluated by Bioanalyzer and qPCR. Sequencing was performed on Illumina HiSeq 2500 or 4000 for single-end 50 bp length reads. Data analysis was as previously described (*34, 38*). Sequence data were mapped against the mm10 genome using BowTie2 (*39*) and signals were normalized by the total numbers of aligned reads and visualized by the UCSC Genome Browser. Peaks were detected by MACS (*40*) with a cutoff of peak length >100 and $-\log_{10}$ (P values) > 50. Peaks of the same mark were merged across CV and/or GF samples from all time points. Spatial analysis between different peaks and genomic features were performed using "countOverlaps" function in R. Circadian analysis was performed by first counting read intensities at peaks. Circadian characteristics were further calculated using JTK. Heat maps were generated by first counting reads across consecutive 50 bp bins across each peak and plotting the values using "heatmap.2" function in R. DNase sensitivity (GSE57919), HDAC3 and ERRα ChIP-seq (GSE63964) data were downloaded from GEO and analyzed similarly as above.

**16*S* rRNA sequencing and data analysis**

Fecal DNA was purified from freshly collected feces using the FastDNA Spin Kit (MP Biomedicals 116560-200) and a FastPrep-24 5G Homogenizer. Sequencing libraries were prepared using the HotStarTaq Plus Master Mix Kit (Qiagen) with primers flanking variable regions V3-V4. Sequencing was performed on a MiSeq following the manufacturer's guidelines. Operational taxonomic units (OTUs) were defined by clustering at 3% divergence (97% similarity). Final OTUs were taxonomically classified using BLASTn against a curated database derived from RDPII and NCBI. Principle component analysis was performed using the "prcomp" function in R and differential abundance analysis was performed using DESeq2 (*41*).

**Table S1: Primer sequences**

| Primer | Sequence |
|---|---|
| Hdac3RTF | CACCAAGAGCCTTGATGCCTT |
| Hdac3RTR | GCAGCTCCAGGATACCAATTACT |
| Cd36RTF | TCATATTGTGCTTGCAAATCCAA |
| Cd36RTR | TGTAGATCGGCTTTACCAAAGATG |
| Cd36ChIPPosF | TCATCAAACAGCATGAATCTCC |
| Cd36ChIPPosR | TACCTGACAGATGGAAAGCAAA |
| Cd36ChIPCtrF | TCAGATGCTAATTTGTGGTTGG |
| Cd36ChIPCtrR | CCAGAAATAGACCCTTGTGAGC |
| Hdac1RTF | AGTCTGTTACTACTACGACGGG |
| Hdac1RTR | TGAGCAGCAAATTGTGAGTCAT |
| Slc25a45ChIPPosF | CCTAGCAGAGAGAGGCAGAGAC |
| Slc25a45ChIPPosR | GCCTGCCTACTACAGTTTTGCT |
| Slc25a45ChIPCtrF | TTACTTCTCAGGCCTCTTCCAG |
| Slc25a45ChIPCtrR | AACCCGATAACCTCCCACTAAT |
| EnpepChIPCtrF | GAAGCAAAGAGAAAAGGCAAAA |
| EnpepChIPCtrR | GGTGCTGAGCTGTATGTGCTAC |
| EnpepChIPPosF | TTGGCTCAGCGCTATATAAACA |
| EnpepChIPPosR | TTTGCCAAGTGATTTCTCTGAA |
| Epas1ChIPPosF | AAAGCAGAAATATTGGGACTCG |
| Epas1ChIPPosR | CAAGGAGTCTGTGTGAACTCCA |
| Epas1ChIPCtrF | AAAAGTGAGGCTGAAAGGACAC |
| Epas1ChIPCtrR | AACTGCGACTTGTTTTGAGGT |

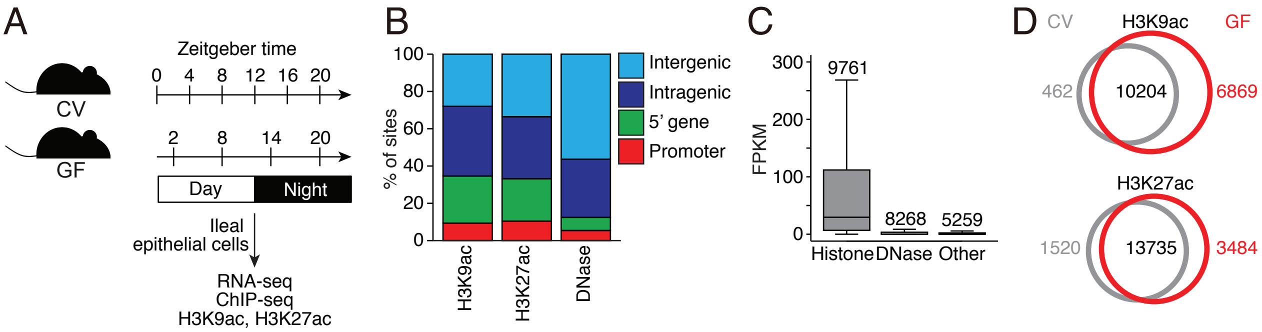


**Figure S1. Chromatin analysis of intestinal epithelial cells from conventional and germ-free mice. (A)** Strategy for chromatin landscape characterization and gene expression analysis. **(B)** Percentages of marks located at promoters, the 5' end of genes, intragenic and intergenic regions. **(C)** Expression levels of transcripts associated with both histone and DNase marks (Histone), DNase mark only (DNase) and neither histone nor DNase marks (Other). **(D)** Venn diagrams showing the spatial similarity of histone acetylation peaks in intestinal epithelial cells from CV and GF mice. Numbers of unique and common peaks between CV and GF samples are indicated. CV, conventional; GF, germ-free; FPKM, Fragments Per Kilobase of transcript per Million mapped reads.

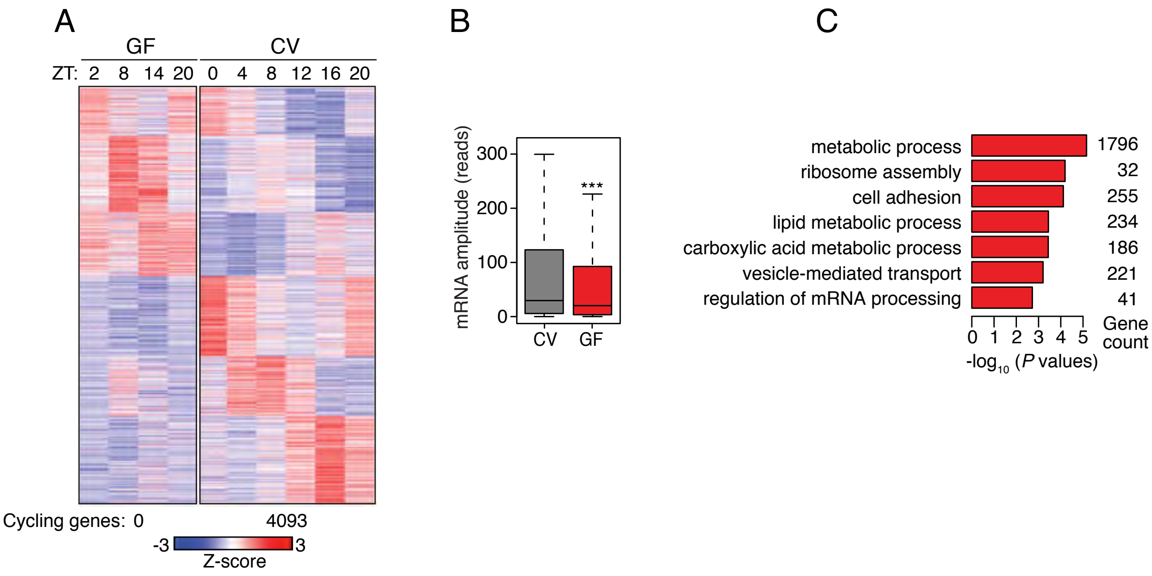


**Figure S2. Transcriptome analysis of intestinal epithelial cells from conventional and germ-free mice.** **(A)** Heat map shows the expression pattern for cycling transcripts across a 24-hour cycle in GF and CV (*7*) epithelial cells (3 pooled biological replicates per library). Rhythmicity was calculated by JTK and the numbers of cycling transcripts are shown at the bottom of the heat map ($P < 0.05$). **(B)** Amplitudes of oscillating transcripts from CV and GF mice. ***, $P<0.001$. **(C)** Enriched Gene Ontology categories among cycling transcripts. CV, conventional; GF, germ-free; ZT, Zeitgeber time.

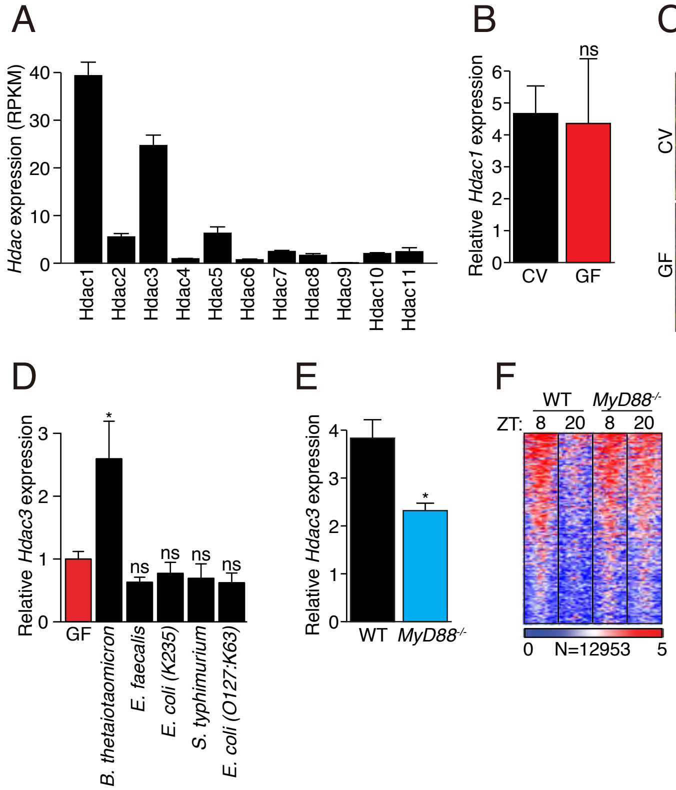


**Figure S3. Analysis of *Hdac* expression in intestinal epithelial cells. (A)** Expression of *Hdac* transcripts in intestinal epithelial cells from conventional mice was assessed by RNA-seq (*7*). **(B)** qRT-PCR analysis of *Hdac1* transcript abundances in CV and GF mice. **(C)** Immunofluorescence detection of HDAC3 in mouse small intestine. Scale bars=50 µm. **(D)** qRT-PCR analysis of *Hdac3* transcript abundance in epithelial cells from GF mice and GF mice after monocolonization with *Bacteroides thetaiotaomicron*, *Enterococcus faecalis*, *Salmonella typhimurium*, the non-pathogenic *Escherichia coli* strain K235 or the pathogenic *E. coli* strain O127:K63. N=3 mice per group. **(E)** qRT-PCR analysis of *Hdac3* expression in epithelial cells from WT and *Myd88*$^{-/-}$ mice. N=3 mice per group. **(F)** Heat map of H3K9ac signals (log (reads at 50 bp windows)) from -1 to +1 kb surrounding the centers of all cycling H3K9ac peaks. N=3 mice per group; *, $P$<0.05 by two-tailed $t$-test; ns, not significant. Means±SEM (error bars) are plotted. RPKM, reads per kilobase, per million mapped reads; CV, conventional; GF, germ-free; WT, wild-type; ZT, Zeitgeber time.

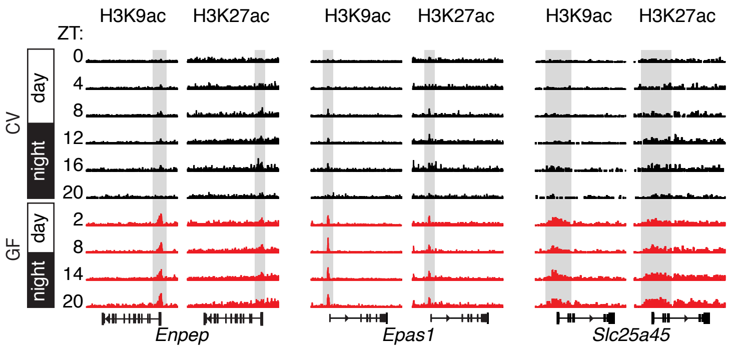


**Figure S4. Histone acetylation at *Enpep*, *Epas1*, and *Slc25a45* in small intestinal epithelial cells exhibits diurnal rhythmicity that depends on the microbiota.** Genome browser view of each gene, showing ChIP-seq analysis of H3K9ac and H3K27ac marks (gray highlights) in small intestinal epithelial cells. The analysis was done across a circadian cycle in CV and GF mice. Each track represents the normalized ChIP-seq read coverage at a single time point. N=3 pooled biological replicates per library. CV, conventional; GF, germ-free; ZT, Zeitgeber time.

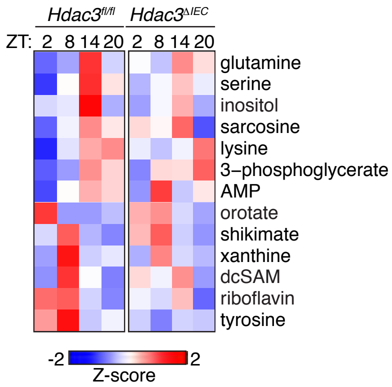


**Figure S5. *Hdac3*$^{\Delta IEC}$ mice show altered diurnal rhythms in serum metabolites.** Serum metabolites were quantified by liquid chromatography and tandem mass spectrometry across a 24-hour day-night cycle and the data are represented as a heat map. Each value in the heat map is the average of three measurements from 3 mice per group. ZT, Zeitgeber time.

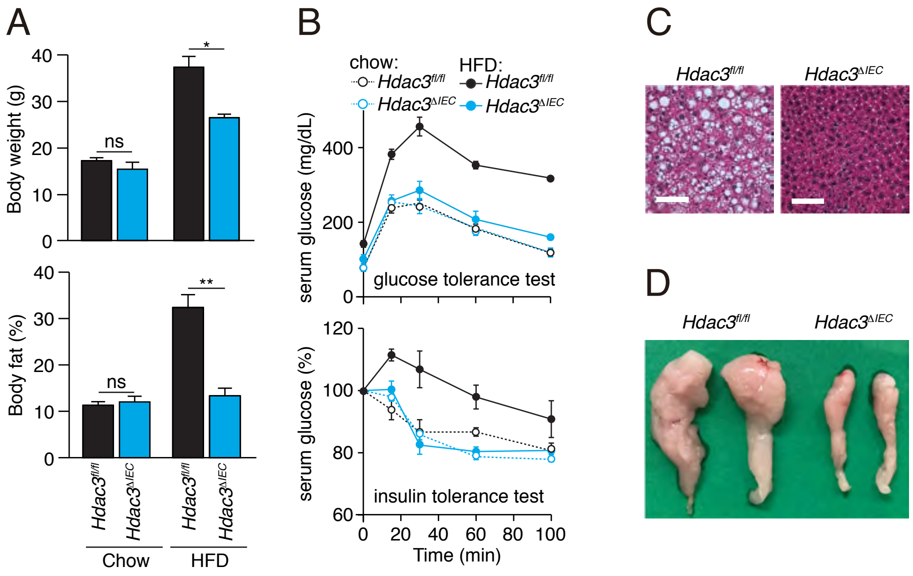


**Figure S6. Metabolic phenotypes of *Hdac3*$^{fl/fl}$ and *Hdac3*$^{\Delta IEC}$ mice. (A)** Weight and body fat percentages of *Hdac3*$^{fl/fl}$ and *Hdac3*$^{\Delta IEC}$ mice fed a chow diet or a high fat diet (HFD) for 10 weeks. N=5 mice per group. *$P$<0.05; **$P$<0.01; ns, not significant by two-tailed $t$-test. Means±SEM (error bars) are plotted. **(B)** Glucose tolerance and insulin tolerance tests in *Hdac3*$^{fl/fl}$ and *Hdac3*$^{\Delta IEC}$ mice fed a chow or a HFD. N=3 mice per group. **(C)** Hematoxylin & eosin staining of liver from *Hdac3*$^{fl/fl}$ and *Hdac3*$^{\Delta IEC}$ mice fed a HFD for 10 weeks. Scale bars=100 µm. **(D)** Epididymal fat pads of *Hdac3*$^{fl/fl}$ and *Hdac3*$^{\Delta IEC}$ mice fed a HFD for 10 weeks. Pictures are representative of three replicates.

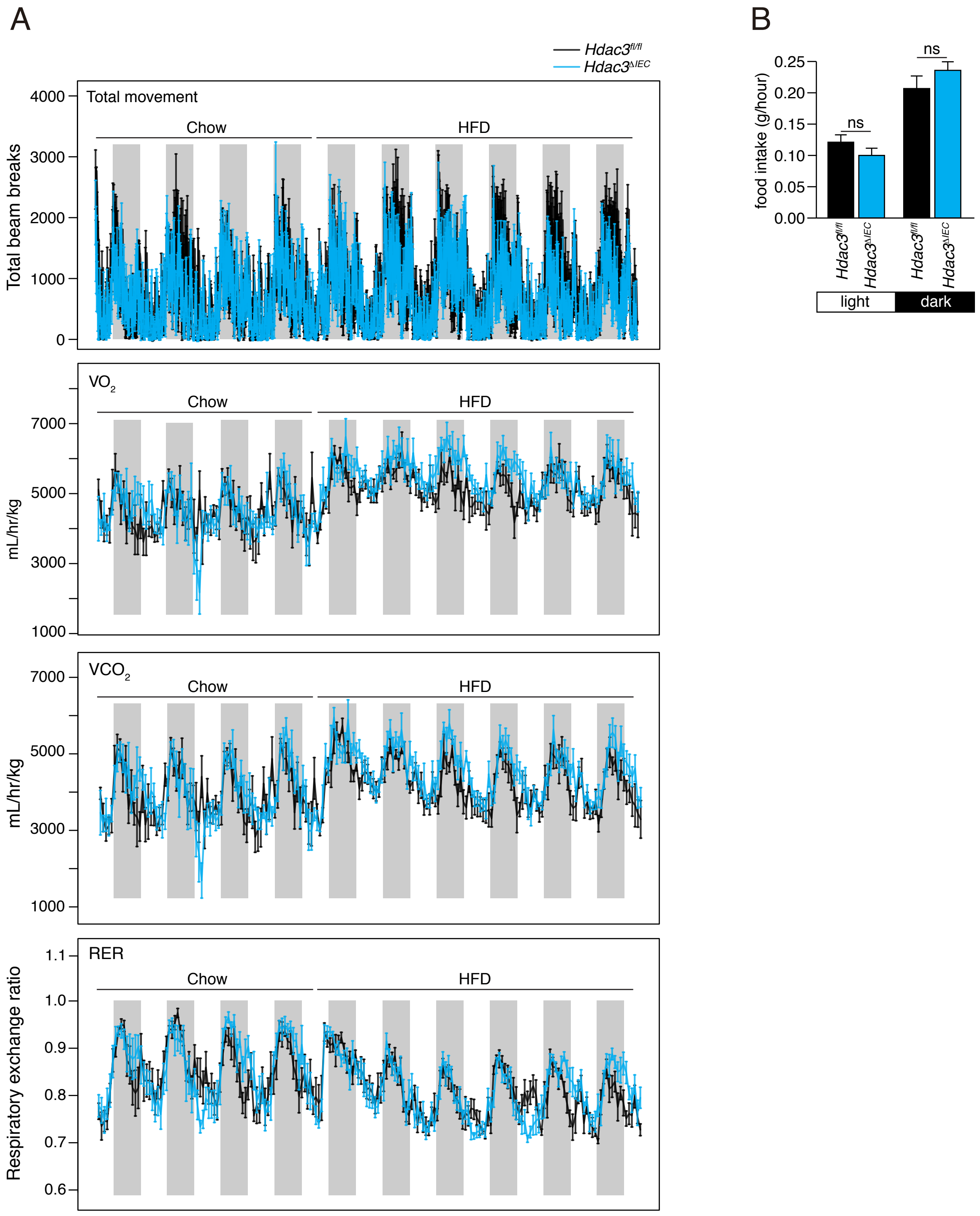


**Figure S7. Physical activity, energy utilization and food uptake are similar in *Hdac3$^{fl/fl}$* and *Hdac3$^{\Delta IEC}$* mice. (A)** Total movement, oxygen consumption rate, $CO_2$ production rate and respiratory exchange ratio (RER) of mice was recorded over 10 days. **(B)** Weight of food eaten by *Hdac3$^{fl/fl}$* and *Hdac3$^{\Delta IEC}$* mice during the daytime and nighttime. All mice were fed a chow diet during the first 4 days and a HFD over the following days. N=6 mice per group. ns, not significant by two-tailed *t*-test. Error bars represent SEM.

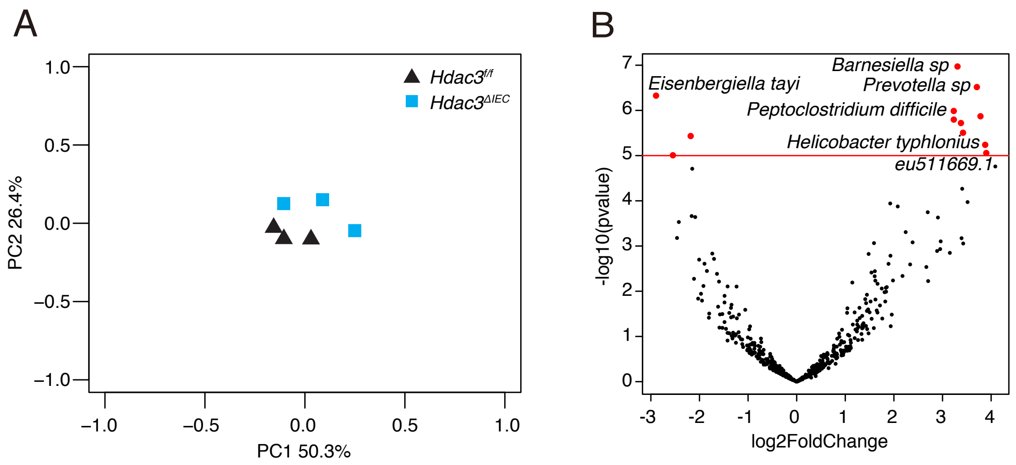


**Figure S8. *Hdac3^fl/fl^* and *Hdac3^ΔIEC^* mice share similar intestinal microbiotas. (A)** Principal coordinate analysis of 16*S* rRNA sequencing of fecal samples from *Hdac3^fl/fl^ and Hdac3^ΔIEC^* mice. The mice were littermates of heterozygous crosses that remained cohoused. Each dot represents one mouse. **(B)** Volcano plot showing bacterial species with differential abundances in *Hdac3^fl/fl^ and Hdac3^ΔIEC^* mice. N=3 mice per group.

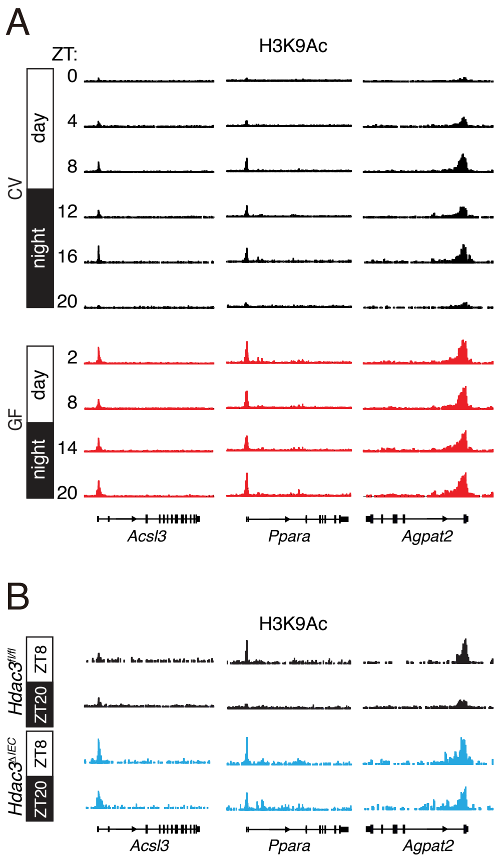


**Figure S9. Histone acetylation at *Acsl3*, *Ppara*, and *Agpat2* in small intestinal epithelial cells exhibits diurnal rhythmicity that depends on the microbiota and *Hdac3*.** Genome browser view of each gene showing ChIP-seq analysis of H3K9ac marks in small intestinal epithelial cells. The analysis was done across a circadian cycle in CV and germ-free GF mice, and at ZT8 and ZT20 in *Hdac3*$^{fl/fl}$ and *Hdac3*$^{\Delta IEC}$ mice. Each track represents the normalized ChIP-seq read coverage at a single time point. N=3 pooled biological replicates per library. CV, conventional; GF, germ-free; ZT, Zeitgeber time.

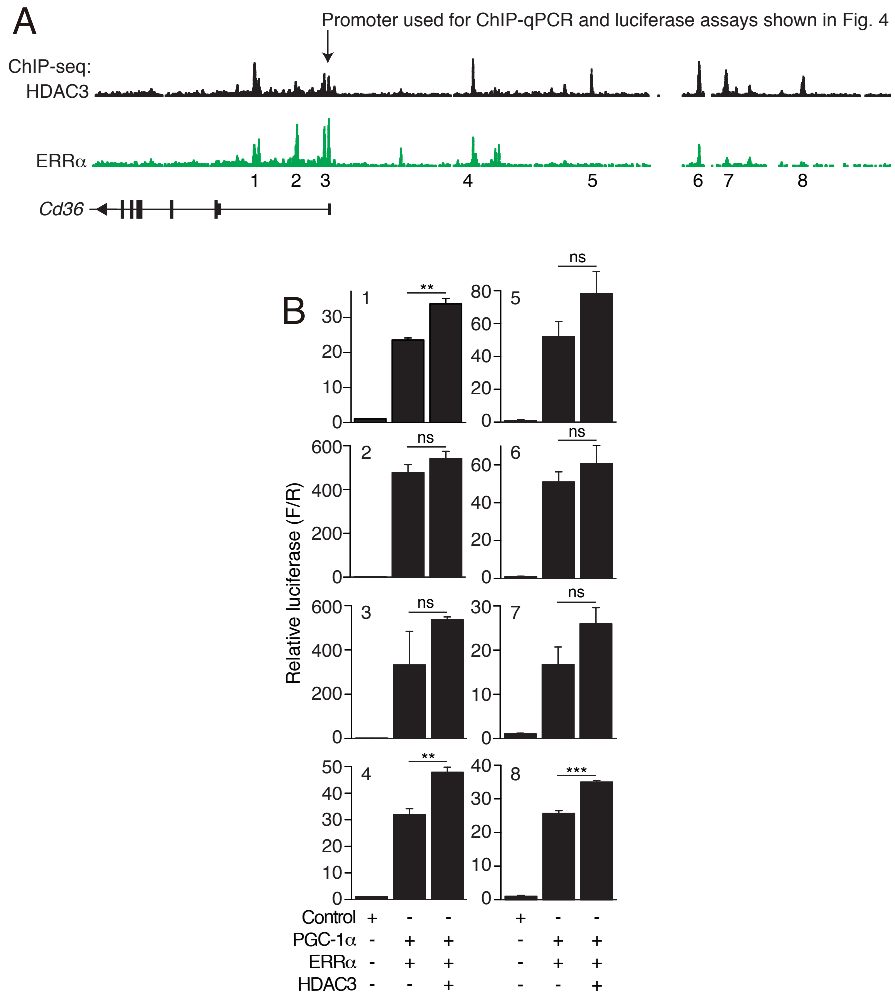


**Figure S10. Transcriptional activation of *Cd36* by ERRα, PGC-1α and HDAC3. (A)** Re-analysis of HDAC3 and ERRα ChIP-seq data collected from brown adipose tissue (*16*) shows that HDAC3 and ERRα colocalize at the *Cd36* promoter and enhancers. **(B)** Luciferase reporter assay of transcription driven by *Cd36* enhancers, demonstrating combinatorial effects of HDAC3, PGC-1α and ERRα. N=3 mice per group. $^{**}P<0.01$; $^{***}P<0.001$; ns, not significant by two-tailed *t*-test; error bars represent SEM.
